# Comparative Restriction Enzyme Analysis of Methylation (CREAM) Reveals Methylome Variability Within a Clonal *In Vitro* Cannabis Population

**DOI:** 10.1101/2023.08.18.552785

**Authors:** Justin Boissinot, Kristian Adamek, Andrew Maxwell Phineas Jones, Eric Normandeau, Brian Boyle, Davoud Torkamaneh

## Abstract

The primary focus of medicinal cannabis research is to ensure the stability of cannabis lines for consistent administration of chemically consistent products to patients. In recent years, tissue culture has emerged as a valuable technique for genetic preservation and rapid production of cannabis clones. However, there is concern that the physical and chemical conditions of the growing media can induce somaclonal variation, potentially impacting the viability and uniformity of clones. To address this concern, we developed Comparative Restriction Enzyme Analysis of Methylation (CREAM), a novel method to assess DNA methylation patterns and used it to assess a population of 78 cannabis clones maintained in tissue culture. Through bioinformatics analysis of the methylome, we successfully detected 2,272 polymorphic methylated regions among the clones. Remarkably, our results demonstrated that DNA methylation patterns were preserved across subcultures within the clonal population, allowing us to distinguish between two subsets of clonal lines used in this study. These findings significantly contribute to our understanding of the epigenetic variability within clonal lines in medicinal cannabis produced through tissue culture techniques. This knowledge is crucial for understanding the effects of tissue culture on DNA methylation and ensuring the consistency and reliability of medicinal cannabis products with therapeutic properties. Additionally, the CREAM method is a fast and affordable technology to get a first glimpse at methylation in a biological system. It offers a valuable tool for studying epigenetic variation in other plant species, thereby facilitating broader applications in plant biotechnology and crop improvement.

## 1. Introduction

Cannabis (*Cannabis sativa* L.) is one the oldest domesticated plants and has a significant economical and societal impact (Torkamaneh & Jones, 2022). It possesses a long history of human use for fiber, oil, seed, and its medicinal and psychoactive properties (Bonini et al., 2018; Russo et al., 2008). Cannabis is a predominantly dioecious diploid annual herbaceous plant (2*n* = 20) that can accumulate a high quantity of specialized phytocannabinoids within its glandular trichomes (Andre et al., 2016). It is known to produce over 545 potentially bioactive secondary metabolites, including more than 177 cannabinoids, various flavonoids, and a plethora of terpenes (Hanuš & Hod, 2020). Despite a large diversity of metabolites produced, the species is often divided and regulated based on the level of a single psychoactive cannabinoid, Δ^9^-tetrahydrocannabinol (THC). In most countries (e.g., Canada, the U.S.A., the E.U.), plants that produce less than 0.3% THC are regulated as hemp, while plants producing 0.3% or more are classified as drug-type. In 2022, the global legal drug-type cannabis market was valued at USD 27.7 billion and is projected to reach USD 82.3 billion by 2027 (Markets and Markets, August 2022, https://www.marketsandmarkets.com/Market-Reports/cannabis-market-201768301.html). Despite the rapid commercial growth of this crop, its biology remains poorly understood due to its long history of prohibition.

Although cannabis is widely used for medicinal and recreational purposes, there are concerns about the consistency and reproducibility of the derived products. This variation is due to a combination of each plant’s genome, as well as the environment in which it is grown, referred to as genotype by environment (GxE) interactions (Booth et al., 2017; Campbell et al., 2019). Within the genomic component, there can be genetic mutations as well as epigenetic differences that can both contribute to differential phenotypic expression. In order to reliably produce consistent extracts, it is critical that they are obtained from genetically stable plants grown under highly controlled conditions. Although cannabis is an outcrossing species with exceptionally high levels of within population variability, clonal propagation methods are relatively easy to use and are optimized to produce uniform populations (Monthony et al., 2021). As a result, in recent years, clonal propagation methods have emerged as the primary method for large-scale production of cannabis. These methods include taking cuttings from selected mother plants (i.e., specific plants with desirable growth characteristics and chemical composition that are maintained in a vegetative stage or in tissue culture for extended period of time), ensuring their proper rooting and growth under controlled conditions, and utilizing specialized cloning media or plant growth regulators to stimulate root development (Adamek et al., 2022). By carefully implementing these methods, growers can achieve consistent and uniform cannabis populations for mass production. However, anecdotal reports indicate that clonal lines tend to decline in quality over time, leading to clones with reduced vigor and lower levels of cannabinoids compared to the original mother plant. A recent study documented a significant amount of intra-plant genetic diversity within a mother plant (Adamek et al., 2022). This diversity could impact the long-term genetic fidelity of clonal lines.

An alternative approach to clonal propagation is micropropagation using plant tissue culture techniques to mass-produce plants in a controlled environment. The compact setup of *in vitro* tissue culture allows for a higher density of plants, minimizing the floor area needed for maintaining mother plants. Importantly, the sterile nature of this technique enables the production of insect-, pathogen-, and virus-free propagules, reducing biotic pressures on the plants (Hesami et al., 2021; Monthony et al., 2021). Typically, it is expected that clones produced *in vitro* using tissue culture techniques will share the same genetics and thus express the same phenotypes. However, somaclonal variations, i.e., genetic or epigenetic induced phenotypic variations between clones produced in tissue culture, are extensively reported in the literature (Bairu et al., 2011; Larkin & Scowcroft, 1981). Epigenetic regulation has been identified as a major cause of these variations since it affects the gene expression of seedlings at different growth and developmental stages (Bednarek & Orłowska, 2020; Miguel & Marum, 2011). Epigenetic factors are heritable and potentially reversible modifications who influence gene expression without altering the DNA sequence. They include processes such as histone state modifications, noncoding RNAs and DNA methylation, which collectively influence chromatin structure (Lauria & Rossi, 2011). They have been hypothesized to be linked to rejuvenation in several plant species (Z. Zhang et al., 2020), including cannabis (Hesami et al., 2023). Among these factors, DNA methylation is widely studied and prevalent in plants. It involves the addition of a methyl group to specific cytosine residues in different contexts (i.e., CG, CHG and CHH) (Springer & Schmitz, 2017). Recent studies documented that in a tissue culture setting, modification of DNA methylation patterns in the genome is more common and is associated with changes in DNA sequence, chromosome breaks and activation of transposable elements (TEs) that can influence gene regulation, chromatin inactivation and cell differentiation (Ghosh et al., 2021).

To date, different methods have been developed and applied in different species to profile the methylation landscape across genomes (Yong et al., 2016). These methods vary in DNA input, resolution, genomic region coverage and bioinformatics analysis (Bock, 2012). Currently, selecting a suitable approach requires an in-depth knowledge of these methods. Despite significant decrease in sequencing costs and advances in bioinformatics analysis, whole-genome methylome profiling remains expensive in the context of large-scale studies. Hence, different low-cost approaches such as microarray-based DNA methylation profiling techniques, restriction enzyme-based and reduced representation bisulfite sequencing (RRBS) methods were developed and widely used for detecting methylated regions (thoroughly reviewed by S. Li & Tollefsbol, 2021). Regardless, their application in large populations remains limited. Moreover, despite analyses of the patterns and effects of DNA methylation in plants (Ghosh et al., 2021; H. Zhang et al., 2018), questions such as the accumulation and location of epimutation sites remain unresolved (Hazarika et al., 2022; Us-Camas et al., 2014).

In the context of cannabis production, determining whether clonal lines derived from tissue culture are uniform is crucial for the consistency and reproducibility of the products. In addition, it is essential for maintaining and preserving germplasm, elite genotypes or parental lines used in breeding programs (Adhikary et al., 2021). Very strict and rigorous quality control and assurance processes as well as the standards related to the safety of cannabis products for medicinal applications require the most precise and regulated production chain (MacCallum et al., 2022; Pusiak et al., 2021). Furthermore, product quality depends on agronomic and environmental factors during plant growth, but also inevitably on the genetic and epigenetic fidelity of the cultivated varieties (Backer et al., 2019). Since micropropagation of uniform clonal lines via tissue culture is fundamental to the cannabis industry, it is thereby critical to study the genetic and epigenetic variations of plants to ensure their long-term stability. The concept of epi/genetic uniformity (or fidelity, stability) can be defined as the absence of variation in the epigenome (epigenetic) and the DNA nucleotide sequences (genetic) within clonal lines.

In this study, we developed a fast and affordable methylotyping method, the Comparative Restriction Enzyme Analysis of Methylation (CREAM), to assess DNA methylation patterns. The CREAM approach, coupled with our bioinformatics pipeline, enabled us to evaluate DNA methylation in a population of 78 cannabis plants representing two clonal lines maintained *in vitro*. This study not only introduces a highly efficient and reliable tool for identifying methylated regions but also provides valuable insights into the methylome uniformity of clonal lines derived from *in vitro* tissue culture.

## 2. Methods

### 2.1 Plant material and DNA extraction

The cannabis clonal population used in this study was initiated in March 2019 at the University of Guelph (Ontario, Canada). It was developed using two sister lines (seedlings) derived from a cross of a cultivar exhibiting an *indica*-leaning growth habit and THC and cannabidiol (CBD) levels of approximately 13% and < 1%, respectively (Adamek et al., in press). Nodal explants from the seedlings were subcultured *in vitro* and maintained on DKW Basal Medium with Vitamins (Product ID D2470; Phytotechnology Laboratories, Lenexa, Kansas, USA), 1 mL/L plant preservative mixture (PPM; Plant Cell Technology, Washington, DC, USA), 0.6% agar (w/v) (A360-500; Fisher Scientific, Fair Lawn, New Jersey, USA) and pH adjusted to 5.7 to generate two clonal lines (Page et al., 2021). A total of 78 clones (from this existing population) of the same chronological age but maintained with different subculture frequencies (i.e., number of subcultures ranging from 6 to 11) were selected (Supplementary Figure 1). DNA samples from the 78 clones were extracted from plant stem cells collected at the same time and from the same stem regions using a Qiagen DNeasy Plant Mini Kit according to the manufacturer’s protocol. The DNA concentration of each sample was quantified using a NanoDrop One spectrophotometer (Thermo Scientific, Waltham, MA) and then diluted to 10 ng/μl. A volume of 10 μl containing 100 ng of DNA was used to prepare each sequencing library for each sample.

### 2.2 CREAM libraries preparation

Two sequencing libraries were prepared in parallel for each sample with the extracted DNA using the Comparative Restriction Enzyme Analysis of Methylation (CREAM) approach (Figure 1) at the Plateforme d’Analyses Génomiques (http://www.ibis.ulaval.ca/en/services-2/genomic-analysis-platform/) at the Institut de Biologie Intégrative et des Systèmes (IBIS) of Université Laval, Quebec, Canada. The CREAM method builds on the 3D-GBS approach (de Ronne et al., 2023). Briefly, in both libraries, DNA molecules were cleaved at one end either by the restriction enzymes *Nsi*I or *Pst*I, which have distinct restriction sites (5’-ATGCA/T-3’ and 5’-CTGCA/G-3’, respectively) to anchor DNA fragments at specific locations in the genome. Since they recognize specific sequences of six nucleotides (six-cutters) and each nucleotide has one of the four nucleic bases, these enzymes cut at a theoretical frequency of 4^6^ or 4096 base pairs (bp). The combined use of the enzymes reduces this theoretical frequency by half, anchoring the DNA fragments at every 2048 bp and thus covering a larger part of the genome. The actual frequency of restriction sites varies between species and within a single plant genome and is mostly influenced by the percentage of GC bases (Torkamaneh et al., 2021). The other end of the DNA fragments was cleaved by either *Msp*I or *Hpa*II, depending on the library. These enzymes share the same restriction site (5’-C/CGG-3’) but have different sensitivity to DNA methylation on the second cytosine (at the CpG site), *Msp*I being insensitive and *Hpa*II being sensitive to methylation. As illustrated in Figure 1a, the variation in sensitivity of the restriction enzymes to DNA methylation results in the generation of fragments of different lengths within the same genomic region. All four possibilities expected from this comparative analysis of the restriction fragments in both libraries, based on the filtering of the fragments with the size selection step, are represented in Figure 1b. After digestion, the sample-specific barcodes and universal adapters were ligated. Since the cohesive ends for the *Msp*I and *Hpa*II restriction sites are identical, the same adapters were used in both libraries. A size selection step using a BluePippin apparatus (Sage Science, Beverley, MA, USA) was performed to capture digested fragments of 200-400 bp. Finally, DNA libraries were amplified by PCR and sequenced with an Illumina NovaSeq 6000 System at the Centre d’expertise et de services Génome Québec (Montreal, QC, Canada), generating 204 M and 173 M paired-end reads of 150 bp for the *Msp*I and *Hpa*II libraries, respectively.

**Figure 1.**
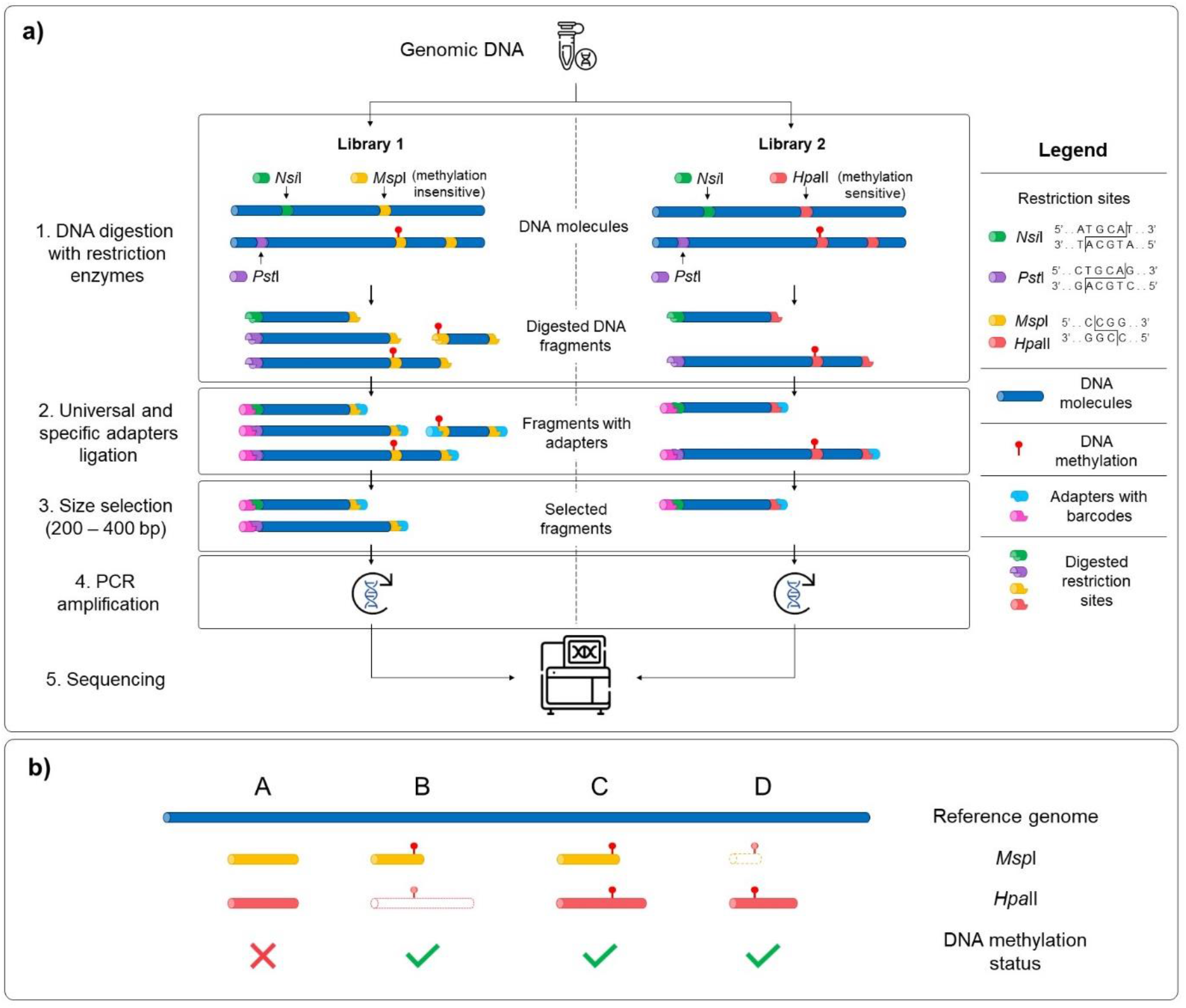
Schematic representation of the Comparative Restriction Enzyme Analysis of Methylation (CREAM) approach. **a)** The same genomic DNA was used as input for the preparation of two libraries. Digestion of the DNA molecules was performed using a set of restriction enzymes sharing a same restriction site but with different sensitivity to DNA methylation (1). Then, universal adapters and sample-specific barcodes were ligated to digested fragments (2). DNA fragments were size selected (3), amplified (4) and sequenced (5). **B)** The comparative analysis of sequencing data of both libraries leads to four possibilities based on either presence/absence or the length of the DNA fragments. Shared fragments in both libraries with the same length and location in the genome indicate the absence of DNA methylation (possibility A) while differences in the presence or the length of the fragments between the libraries indicate the existence of DNA methylation (possibilities B, C and D).

### 2.3 Bioinformatics pipeline

#### 2.3.1 Alignment of reads

The paired-end sequencing reads were demultiplexed using Sabre (https://github.com/najoshi/sabre) and trimmed (i.e., removing adapters) with cutadapt (Martin, 2011). Then, they were aligned to the cannabis reference genome (cs10 v2 (GenBank Accession No. GCA_900626175.2); (Grassa et al., 2021) with BWA-MEM (H. Li, 2013). Only reads with a high mapping quality (MAPQ score >= 20) were retained for methylotyping. Genome coverage and depth of coverage were obtained using the bedtools genomecov command (Quinlan & Hall, 2010) and samtools coverage command (Danecek et al., 2021).

#### 2.3.2 Methylotyping

Four possibilities can be expected for the mapped reads (Figure 1b). First, there is no DNA methylation if the fragments are of the same length and mapped to the same location of the genome in both (*Msp*I and *Hpa*II) libraries (possibility A). The remaining possibilities capture DNA methylation if fragments are found in only one library (possibilities B or D) or if fragments of different lengths are observed (possibility C). To capture these possibilities, a custom pipeline (https://github.com/justinboissinot/CREAM) programmed in Python 3 (https://www.python.org/) was developed and used to determine methylated regions. This pipeline takes the alignments (BAM files) as an input and outputs the methylated and unmethylated positions across the genome. A brief description of the different steps implemented in this pipeline are provided below.

The paired-end reads from the BAM files that are accurately mapped to their corresponding pair were used to reconstruct the insert fragment from which they originated. This step ensured that both restriction sites were present in the insert fragments, as DNA methylation can occur on only one side of the restriction fragments. Various quality metrics, including the number of differences between the sequence and the reference (distance), the length of CIGAR strings (indicating insertions or deletions in the sequence), and the number of mismatches in the alignment (alignment score) were extracted from the information generated by BWA-MEM. Subsequently, a table of inserts was generated for each sequencing library, which was further utilized for downstream analysis. Then, from both libraries, a list of loci was extracted based on the inserts. In this context, a locus refers to a genomic region that includes the leftmost and rightmost positions of a set of inserts that either overlap between two libraries or are non-overlapping in either library (Supplementary Figure 2). Loci with overlapping inserts represent regions where the two libraries share at least one nucleotide overlap, including inserts that share the same restriction site or have perfectly overlapping inserts (from their leftmost position to their rightmost position). Loci with non-overlapping inserts represent regions where an insert is present in only one library, capturing regions unique to each library in the loci list. A locus was excluded from the list if less than half of the samples had a coverage of under 20X (inserts) for that specific locus. For each locus, a methylation status was determined based on the aforementioned possibilities (Figure 1b). Methylated and unmethylated positions from the loci were then separated and saved in different files for subsequent analysis.

### 2.4 Accumulation and distribution of methylated positions

To examine the accumulation of methylated loci in the population as the number of subcultures increased, a Kruskal-Wallis test (one-way ANOVA on ranks) was conducted. This test was chosen since the assumptions for conducting an ANOVA were not met in this case. Then, a principal component analysis (PCA) was performed to assess the distribution of methylated positions within the population. The PCA aimed to determine clustering patterns in the samples based on the methylated loci using the R packages *FactoMineR* and *factoextra* (Kassambara & Mundt, 2016; Lê et al., 2008).

The distribution of methylated regions across the cannabis genome was visualized using the *RIdeogram* R package (Hao et al., 2020). This also included the gene density information obtained from the NCBI Gene table for the cs10 reference genome (https://www.ncbi.nlm.nih.gov/data-hub/gene/taxon/3483/, accessed February 15, 2023). A Spearman’s rank correlation coefficient was calculated, using the *cor* command from the R package *stats* (R Core Team, 2022), to assess the monotonic relationship between the gene density and the methylation density across the genome since the data for both variables were skewed towards 0. Other visualizations were generated with *ggplot2* in R (Wickham, 2011).

### 2.5 Gene Ontology (GO) analysis

A gene ontology (GO) analysis was performed to identify significant GO terms affected by DNA methylation captured with the CREAM approach. To overcome challenges in matching protein IDs and GO terms with the annotations of the cs10 cannabis reference genome, a combination of the GAWN v0.3.5 (https://github.com/enormandeau/gawn) and go_enrichment (https://github.com/enormandeau/go_enrichment) pipelines was used. The GAWN pipeline annotated the cs10 reference genome using the available transcriptome from NCBI and found all the methylated loci and the captured loci (unmethylated and methylated) within ± 1 kb of transcripts. The GO enrichment analysis was then performed with the go_enrichment pipeline using the list of methylated loci adjacent to transcripts as the target and considering all captured loci as the background gene set. GO terms with a significant adjusted *p*-value of *p* < 0.10 (Benjamini-Hochberg false discovery rate correction for multiple testing) were kept for further exploration of biological processes.

## 3. Results

### 3.1 Development and validation of the CREAM approach

We have developed a low-cost and efficient method for assessing methylome variability at the population level. This method, CREAM, involves digesting DNA samples with restriction enzymes of different sensitivity to DNA methylation. Libraries were developed and sequenced for 78 cannabis clones produced in tissue culture. This has yielded an average of >188 M paired-end reads per library (Table 1). The demultiplexing, trimming and alignment of the reads to the reference genome led to an average of >2 M paired-end reads per sample.

**Table 1.**
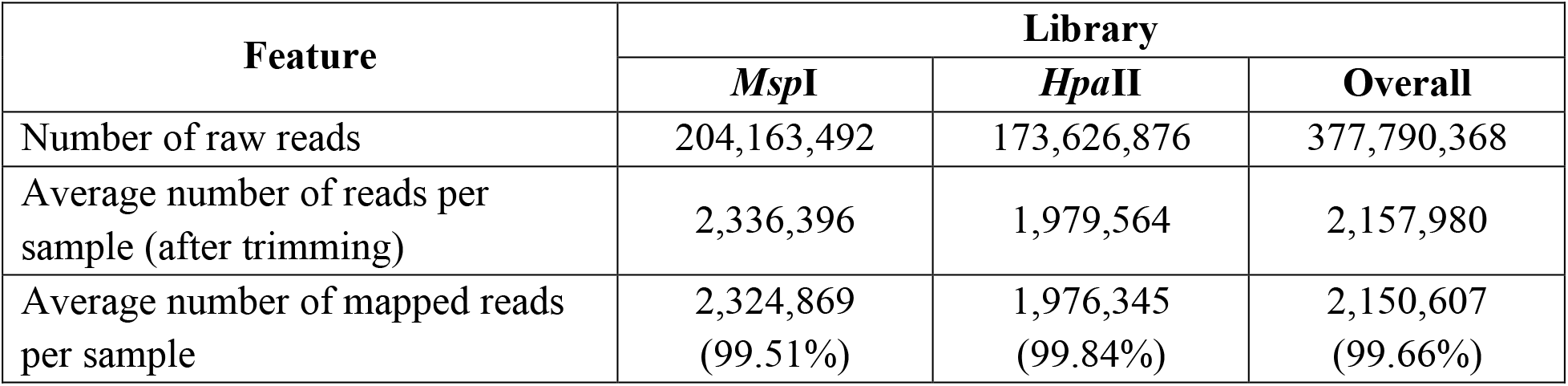
Summary of sequencing libraries statistics.

**Table 2.** List of enriched GO terms that are influenced by DNA methylation in cannabis.

| GO term | Name of biological process | *p*-value (FDR BH) |
| --- | --- | --- |
| GO:0006139 | Nucleobase-containing compound metabolic process | 0.00001 |
| GO:0046483 | Heterocycle metabolic process | 0.00001 |
| GO:0090304 | Nucleic acid metabolic process | 0.00003 |
| GO:0006725 | Cellular aromatic compound metabolic process | 0.0001 |
| GO:1901360 | Organic cyclic compound metabolic process | 0.0002 |
| GO:0034641 | Cellular nitrogen compound metabolic process | 0.0003 |
| GO:0016070 | RNA metabolic process | 0.009 |
| GO:0006807 | Nitrogen compound metabolic process | 0.011 |
| GO:0016071 | mRNA metabolic process | 0.045 |
| GO:0009987 | Cellular process | 0.084 |
| GO:0006325 | Chromatin organization | 0.084 |

Out of the initial 78 samples, 10 samples were excluded from the analysis as they yielded less than 100,000 reads on average per library. The genome coverage and mean depth of coverage were computed for the remaining samples and compiled for each library (Supplementary Table 1). On average, we captured ∼0.4% of the cannabis genome with a mean depth of coverage of ∼100X across the captured regions, indicating a sufficient depth to ensure reliable and accurate analysis of the captured regions. As shown in Figure 2, the genome coverage tends to be higher in the *Msp*I library, while the *Hpa*II library exhibits a higher mean depth of coverage. This disparity can be attributed to the fact that the *Hpa*II library does not capture DNA fragments with DNA methylation. Consequently, the *Hpa*II library contains fewer fragments for a comparable sequencing effort compared to the *Msp*I library, covering smaller proportion of the genome with higher coverage.

**Figure 2.**
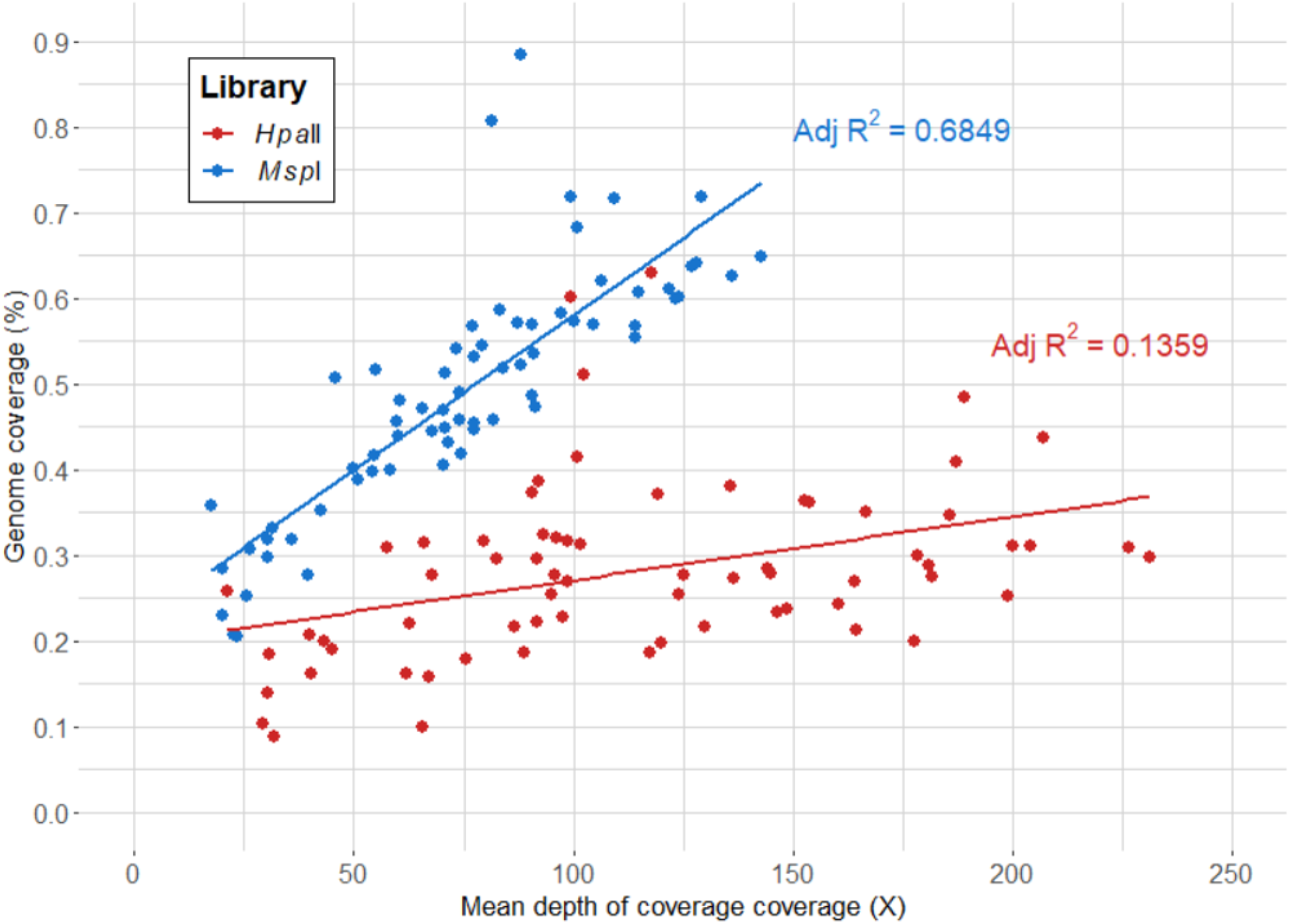
Genome coverage (%) per mean depth of coverage (X) for each library.

### 3.2 Identification of methylated positions

From the mapped paired-end reads, a bioinformatics analytical pipeline was developed to determine the methylated positions captured with the CREAM approach. Briefly, the pipeline includes quality checks for inserts (composed of a pair of reads) and calls loci that encompass all potentially overlapping inserts within a region (See Supplementary Figure 2 for details). This methylotyping pipeline successfully divided the inserts from the CREAM libraries into the four expected possibilities, resulting in a total of 5,235 loci (See Figure 1b for the four expected categories). Of these confidently called high-quality loci, 3,762 (71.86%) were identified as methylated regions, while 1,473 (28.14%) were identified as unmethylated regions with perfectly overlapping inserts in both libraries (possibility A). The most common type of captured methylated regions (2,949) were loci with an insert only in the *Msp*I library (possibility B), accounting for 56.33% of all captured loci. Loci with an insert only in the *Hpa*II library (possibility D) accounted for approximately 15.53% of all captured loci (813). Finally, a total of 186 loci were excluded from the analysis due to challenges in accurately categorizing them into the predetermined possibilities. These included methylated regions composed of loci with inserts of different length in both libraries (possibility C), which were far less common and loci that exhibited ambiguous characteristics or lacked clear patterns for classification, rendering them less reliable for further analysis.

### 3.3 Distribution of methylated positions in the population

The methylotype of each sample for the list of called loci was used to assess the variability in methylated positions across the clonal population. The frequency of the methylated loci revealed four main peaks (Figure 3a). The largest peak consisted of the 1,490 (39.61% of total methylated loci) monomorphic methylated loci, which are DNA methylation positions captured in all 68 samples, indicating a shared methylotype across the population. On the other hand, among polymorphic loci, 427 (11.35%) were unique to a single sample, indicating specific methylation patterns within individual samples. The two other peaks in the distribution of the number of methylated loci corresponded to the number of clones derived from the two sister lines. Specifically, there were 155 loci unique to the 13 samples in the AT-4 line and 117 loci unique to the 55 samples in the AC-150 line, indicating distinct methylotypes within the sister clonal lines.

**Figure 3.**
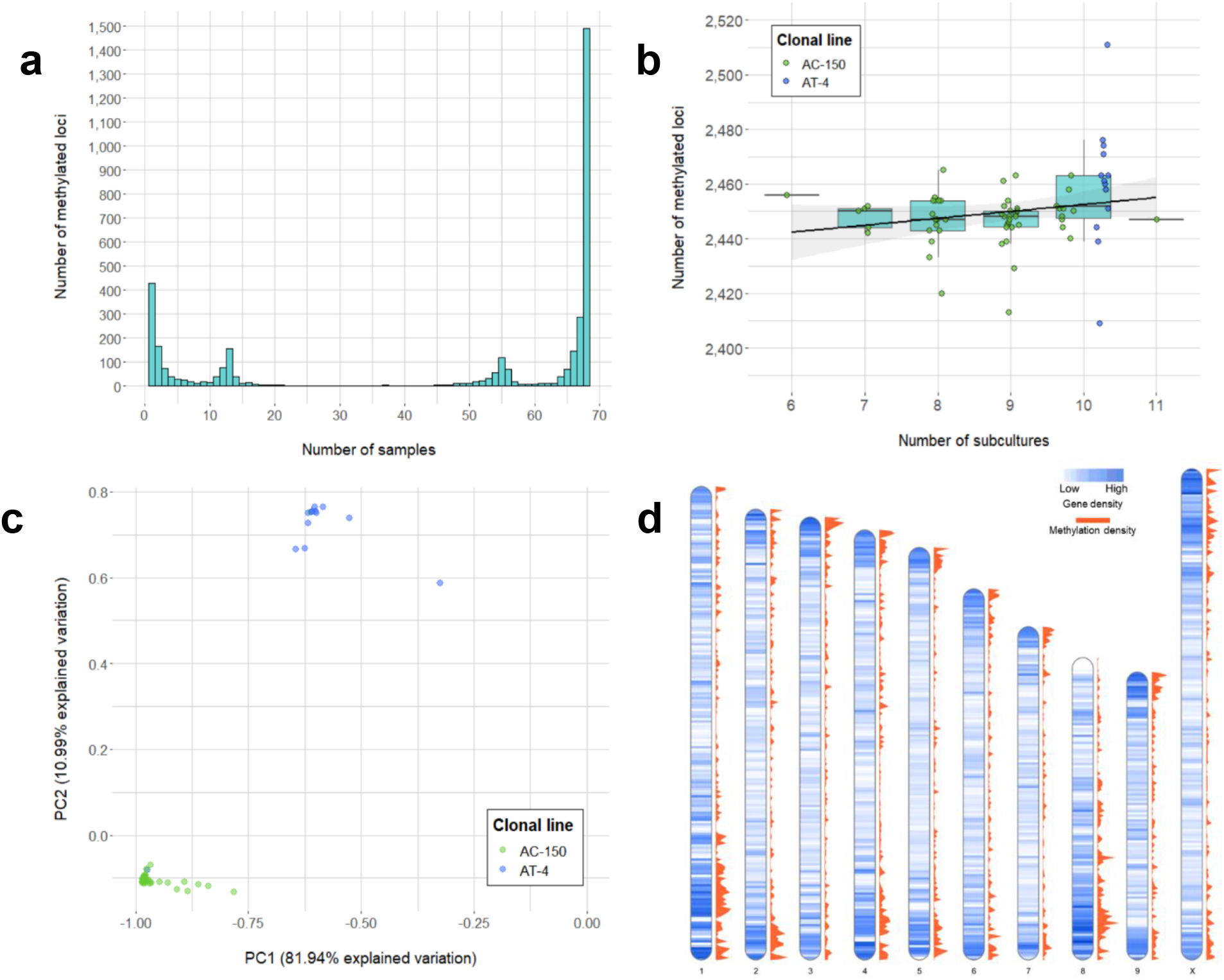
Number of methylated loci called from the CREAM libraries. **a)** Frequency of methylated loci within the population. **B)** Number of methylated loci per number of subcultures for both clonal lines (AC-150 line in green and AT-4 line in blue). **C)** Principal component analysis (PCA) of methylated loci called from the CREAM libraries. Both clonal lines derived from two sister lines are clustered into two different groups. **D)** Density of genes and methylated loci on the chromosomes of the cs10 cannabis reference genome in bins of 500 kb.**Table 1**. Summary of sequencing libraries statistics.

In this study, we also investigated whether the number of subcultures that the clonal lines went through in tissue culture had an impact on the number of DNA methylation positions captured. Although a significant methylome variability within clonal lines was observed, we did not find a significant correlation between the number of subcultures and the total number of methylated loci (*p*-value = 0.19; one-way ANOVA on ranks (Kruskal-Wallis test)) (Figure 3b). This suggests that the variability in DNA methylation within clonal lines was not influenced by the number of subcultures the plants went through in this study. Finally, the principal component analysis (PCA) performed on the methylated loci (Figure 3c) revealed that the two principal components explained a significant portion of the variation, accounting for 81.94% and 10.99% of the total variation, respectively. The samples displayed clear clustering patterns, with two distinct groups corresponding to the clones derived from the two sister lines. Within each cluster, the variability was relatively limited, although some samples showed variation compared to overall trend of their respective group.

### 3.4 Distribution of the methylated positions in the genome

We examined the distribution of the methylated positions across the genome to understand the patterns of DNA methylation. Precisely, we calculated the gene density, which represents the number of genes present in each bin of 500 kb in the cannabis genome, along with the methylation density, which indicates the number of captured methylated positions within the same 500 kb bin (Figure 3d). We then compared the gene density and methylation density to understand the relationship between DNA methylation and gene distribution in different genomic regions. The density of methylated loci across the genome showed a strong positive correlation with the density of genes across the same genomic bins (Spearman’s rank correlation coefficient = 0.6473), suggesting the positive monotonic relationship between the two variables. We also observed that regions of high genic density were also enriched in captured methylated loci. However, the observed correlation between gene density and methylation density could be influenced by the approach used, which tends to capture loci in genic regions (Supplementary Figure 3). Finally, the gene ontology (GO) analysis was performed to identify genes potentially affected by the DNA methylation patterns captured within a range of ± 1 kb of gene transcripts. The analysis utilized all the loci captured by the CREAM approach as a background set for comparison. As a result, 11 GO terms were found to be significant. Most importantly, 9 out of the 11 significant biological process GO terms were related to metabolic processes, indicating a potential influence of DNA methylation on important biochemical pathways.

## 4. Discussion

In cannabis and other plant species produced through micropropagation, it is crucial to understand the underlying factors contributing to somaclonal variation. While there has been extensive research on the influence of media culture conditions and genetic variation (reviewed in Krishna et al., 2016; Sato et al., 2011; D. Zhang et al., 2014), epigenetic factors have recently emerged as noteworthy factors that could account for phenotypic variability that cannot be explained by genetic mutations in micropropagated plants. Of particular interest is the examination of DNA methylation patterns (Bobadilla Landey et al., 2015; Borges et al., 2021; Han et al., 2018; Jaligot et al., 2000; H. Li et al., 2012; Matthes et al., 2001; Ong-Abdullah et al., 2015; Wibowo et al., 2022). Despite an increasing interest in cannabis as a valuable crop, there is a notable gap in research regarding DNA methylation and its influence on the cannabis genome, particularly in the context of tissue culture. This knowledge gap hinders the development of effective strategies to ensure reproducibility and efficiency in tissue culture practices, which are essential for various production systems. Therefore, the primary objective of this study was to develop a novel, low-cost, and high-throughput approach (i.e., CREAM) to detect variations in the methylome of *C. sativa* clones derived from *in vitro* tissue culture. By investigating DNA methylation patterns, we aimed to shed light on epigenetic factors contributing to somaclonal variation in cannabis and provide insights into the potential regulatory role of DNA methylation in shaping the phenotypic diversity observed among clones.

### CREAM: a new efficient and cost-effective methylotyping method

The CREAM method was successful in identifying significant variation in methylotypes among cannabis clones, specifically distinguishing the two subpopulations derived from two sister lines. This success can be attributed to the generation of high-quality sequencing fragments, which facilitated the accurate identification of methylated and unmethylated loci using our newly developed methylotyping pipeline. Importantly, the CREAM method offers a cost-effective solution compared to the main approaches based on RRBS, like epiGBS (Van Gurp et al., 2016) and bsRADseq (Trucchi et al., 2016), with an estimated sequencing cost per sample of around 30$, representing a 30-80% decrease in costs (Werner et al., 2020). Additionally, the CREAM approach avoided the extensive DNA damage typically associated with the bisulfite conversion in other techniques (Tanaka & Okamoto, 2007), preserving DNA integrity.

The CREAM method achieved a satisfactory average genome coverage of ∼0.4% and a mean depth of coverage of around 100X. In comparison, well-regarded RRBS methods such as epiGBS (Van Gurp et al., 2016) and its modified version (Werner et al., 2020) covered 0.37% of the 135 Mb of the *A. thaliana* genome and 0.28% of the 246 Mb almond genome, respectively. When examining the genome coverage and mean depth of coverage specific to each library (Figure 2), it becomes evident that the genome coverage tends to be higher in the *Msp*I library compared to the *Hpa*II library. This observation is consistent with the higher proportion of loci with fragments found only in the *Msp*I library (category B) among the captured regions. Therefore, the differences in genome coverage and the number of loci found between the two libraries can be attributed to the variations in the efficiency of the two restriction enzymes (methylation sensitive and methylation insensitive) used in the CREAM method.

The choice of the restriction enzymes in the CREAM method has a direct impact on the number of loci captured and their distribution across the genome. In our findings, we observed a notable enrichment of fragments within genic regions. To explain the abundance of captured fragments, we propose two potential explanations. The first possibility relates to the influence of GC content as a contributing factor. In the case of the cannabis reference genome (cs10), it exhibits an overall GC content of 33%, whereas the GC content differs among the restriction sites of the restriction enzymes. Specifically, the GC content of the *Nsi*I and *Pst*I restriction sites is 66% and 33%, respectively, while the *Msp*I and *Hpa*II restriction sites have a GC content of 100%. This indicates that the distribution of restriction fragments is not uniform across the genome and tends to be more concentrated in regions that are relatively richer in GC than AT. In plants, these GC-rich regions are predominantly associated with genic regions (Glémin et al., 2014; Serres-Giardi et al., 2012). The second explanation involves the significant role of chromatin structure in the accessibility of enzymes’ restriction sites (Sotelo-Silveira et al., 2018). Typically, plant genomes exhibit distinct compartments with varying accessibility. Euchromatin regions, rich in genes and located at the chromosome tips, are generally more accessible, contributing to the observed concentration of captured fragments within genic regions (Dong et al., 2017, 2020). Although this would require further work that is outside the scope of this study, the efficiency of the restriction enzymes could be associated with the remnants of euchromatin in the extracted DNA in solution. Collectively, these factors support the observed correlation between the gene density and methylation density in the data (Figure 3d). Therefore, the CREAM approach might result in an overrepresentation of genic regions due to their higher accessibility to the restriction enzymes.

While our approach offers several advantages, it is important to acknowledge its limitations. First, its resolution is currently limited. It provides a binary methylation status (either 0 or 1) for a given cytosine in a locus, rather than providing a methylation quantitative score at a base level. Although this binary representation still provides valuable insights, a finer resolution would be desirable to gain a more comprehensive understanding of the methylation landscape across the genome. Second, we encountered challenges in classifying 186 captured loci (3.43% of all loci) into the expected categories (A, B, C or D). The comparative analysis of the libraries was challenging for these specific restriction fragments because of the complexity and diversity of their alignment. Lastly, it is important to note that our approach did not operate at a single-cell level. As a result, the methylation patterns obtained from a single sample represent a mixture of methylation patterns from various cells, introducing some level of heterogeneity. However, these limitations might be overshadowed by the compelling low-cost and high-throughput features of the approach.

### Methylation variation in cannabis clones

Upon analyzing the DNA methylation frequency in the population, we observed contrasting patterns between the two subsets of clonal lines. These patterns can be used as methylation tags or methylation fingerprints, allowing us to clearly differentiate individual clones within their respective group. Notably, the methylation patterns exhibited by each clone strongly reflect their clonal lineage, suggesting a unique and identifiable DNA methylation profile for each clonal line. This is consistent with various studies of methylation patterns in tissue culture. For example, in maize, stable changes in differentially methylated regions (DMRs) were observed between independently regenerated clonal lines, highlighting potential “epigenetic footprints” unique to certain lines and maintained over several generations (Stelpflug et al., 2014). A further study has shown consistent (monomorphic) and rare (polymorphic) DMRs maintained across generations that distinguished tissue culture-derived plants, providing more evidence for specific DNA methylation patterns acting as “tags” (Han et al., 2018). It is likely that these patterns originated from the sister seeds that were used to initiate clonal lines and were preserved to some extent throughout multiple generations of subculturing. In sexual reproduction, such as seed production, various mechanisms exist to limit defective and potentially problematic epigenetic variations from one generation to the next, including resetting of DNA methylation state (Quadrana & Colot, 2016). However, clonal propagation, being asexual, lacks these reset processes and is expected to affect DNA methylation stability across the genome (Ibañez & Quadrana, 2023), leading to potential slight variations in methylation patterns.

Different DNA methylation dynamics have been observed in the vegetative state. For instance, high levels of CHG methylation in *Arabidopsis thaliana* shoot apical meristematic stem cells (Gutzat et al., 2020) and low levels of CHH methylation in species with extensive clonal propagation histories (Niederhuth et al., 2016) have been reported. However, trends regarding CG methylation levels in tissue culture are less conclusive and differ among species, with increasing levels reported in gentian (Fiuk et al., 2010), bush lily (Q.-M. Wang et al., 2012), banana (Peraza-Echeverria et al., 2001) and tomato (Smulders et al., 1995), decreasing levels observed in triticale (Bednarek et al., 2017), barley (X. Li et al., 2007), grapevine (Baránek et al., 2010) and *Freesia* (Gao et al., 2010), and no significant changes detected in pea (Smýkal et al., 2007) or apples (X. Li et al., 2002). Since the cannabis methylome is poorly understood, our results represent an encouraging first report of methylation patterns for this economically important crop in a tissue culture context.

Tissue-specific differences in DNA methylation can also be significant (Lloyd & Lister, 2022). Cells from stem tissues are generally less studied and methylation patterns are less decisive, making comparisons between studies and species more intricate. In a study conducted in hops (*Humulus lupulus* L.), a close relative to *C. sativa* (Kovalchuk et al., 2020), micropropagated stem tissues (branches) were tested for methylation changes (Peredo et al., 2009). The study found that most the methylated loci (56.34%) were monomorphic, meaning they were shared by all 80 clones obtained from two cultivars, while 13.24% of the variation were unique to individual samples (referred to as singletons). These results are consistent with the distribution of methylated loci observed in the present study (Figure 3a), with 41.22% and 11.81% of the total loci being monomorphic and unique to a single sample, respectively. The difference in proportions between the methylation patterns observed in our study and those reported in the methylation-sensitive amplified polymorphism (MSAP) method used in the hops study could be attributed to the higher genomic resolution of our approach.

The lack of significant differences in the number of methylated loci with the number of subcultures may be attributed to the dynamic coordination of processes involved in establishing, maintaining, and removing specific DNA methylation states (H. Zhang et al., 2018). It has been reported that while the total number of methylated positions may not change significantly, their specific identity is likely to vary (H. Zhang et al., 2018). This suggests that the stability of methylation patterns is established in the earlier stages of production. The stability of DNA methylation patterns during subculturing has been observed in other plant species as well. For example, in garlic, methylation patterns were found to stabilize after 6 months of micropropagation (Gimenez et al., 2016). However, it is important to note that the trends in DNA methylation variations are highly species-dependant. Some studies have reported increases in methylation levels during subculturing (Fraga et al., 2016; Rival et al., 2013), while others have observed decreases (Huang et al., 2012; Machczyńska et al., 2014; Xu et al., 2004) over different subculturing durations. There have also been cases where methylation levels initially decrease in early generations and then recover over longer periods of time, as shown in sweet orange after 30 years (X. Wang et al., 2022). Considering that our study is the first investigation of the cannabis methylome, it is plausible that methylation patterns stabilized in our clonal population over several subcultures. All clones were acclimatized to tissue culture and could have reached a steady state by the 6^th^ to 10^th^ subculture. Further research is needed to gain a better understanding of the dynamics and stability of DNA methylation patterns in cannabis and how they may influence the phenotypic characteristics of clonal plants. A focus should be made on the induction phase in tissue culture, where plants are transitioning from regular growth conditions to *in vitro* conditions, which could affect methylation patterns.

### Distribution of methylated loci in the cannabis genome

Upon examining the distribution of DNA methylation in the cannabis genome, we observed a correlation between methylation density and gene density along the chromosomes. As mentioned earlier, this suggests that the captured loci in this study may be biased towards genic regions due to their GC-rich content and accessibility. To a lesser extent, non-genic regions were also captured with the CREAM approach. These genomic regions rich in transposable elements and repetitive DNA sequences have also been associated with higher DNA methylation rates in *A. thaliana*, both in heterochromatin and euchromatin (X. Zhang et al., 2006). It is therefore expected to observe methylated loci in these regions. With the current limited knowledge of the cannabis genome, loci found near genes still represent the most interesting regions to study for their immediate potential functional impact on the plants biological processes.

In this study, the GO analysis within a range of ± 1 kb around gene transcripts resulted in 11 significant GO terms. The effect of gene body methylation (gbM) on gene expression and phenotypic changes is still a subject of debate and not fully understood. However, the association between gbM and gene expression is recognized as important, and it represents a potential source of variation that can be subject to natural selection (Muyle et al., 2022), highlighting the importance of the significant GO terms found in this study. Of these 11 GO terms, 9 are related to metabolic processes, which encompass a range of biochemical reactions involved in the synthesis, breakdown, and transformation of various molecules within the plant. This suggests that DNA methylation may play a regulatory role in modulating the expression and activity of genes involved in key metabolic pathways in cannabis. To gain a deeper understanding of the cannabis methylome, it would be crucial to validate the biological effects of DNA methylation on the transcriptional activity of these genes. Such validation experiments would provide important insights into the functional implications of DNA methylation in cannabis and its potential impact on metabolic processes.

## 5. Conclusion

The present study addresses a critical gap regarding somaclonal variation and epigenetic factors in *C. sativa*. It highlights the importance of DNA methylation in shaping phenotypic diversity among plant clones, particularly in the context of tissue culture practices and the reproducibility and uniformity of plant clones. To the best of our knowledge, this is the first study to investigate the cannabis methylome and the first of its kind in the field of cannabis tissue culture. The primary objective of this study was to develop a novel and cost-effective approach to detect methylation variation in *C. sativa* clones derived from tissue culture. The results revealed significant variation in methylotypes among the cannabis clones, indicating the presence of methylation footprints between clonal lines and offering valuable insights into the epigenetic landscape of this important crop. Importantly, this approach overcomes the cost and technical challenges associated with existing methods and enables the high-throughput analysis of methylome at a population level. By elucidating the dynamics of methylome and its correlation with gene density, this study opens avenues for further investigations on the regulatory role of DNA methylation in key metabolic pathways of cannabis. Functional validation of the observed methylation patterns and their impact on gene expression will undoubtedly contribute to a more comprehensive understanding of the cannabis methylome. This knowledge has implications for crop improvement and the development of sustainable production systems in the cannabis industry.

## Author contributions

KA and AMPJ maintained the clonal population. KA collected tissues and extracted DNA. BB and DT conceptualized the sequencing approach. BB acquired the sequencing data. JB and EN conducted data analysis and conceptualized the CREAM workflow. JB and DT contributed to writing the manuscript. All authors read and improved the manuscript.

## Funding

The authors gratefully acknowledge the support of the Natural Sciences and Engineering Research Council (NSERC) of Canada. JB was supported by a NSERC CGS M scholarship and a FRQNT B1X scholarship. None of the funding bodies were involved in study design, data acquisition, data analysis, interpretation of the results, or manuscript writing.

## Notes

https://github.com/justinboissinot/CREAM

### Competing Interest Statement

The authors have declared no competing interest.

