## Supplementals for "Comparative Restriction Enzyme Analysis of Methylation (CREAM) Reveals Methylome Variability Within a Clonal *In Vitro* Cannabis Population"

**Supplementary Methods**

**Supplementary Figure 1.** Population of cannabis clones used in this study. Clonal lines are represented in green (AC-150) and blue (AT-4). Population tree was loaded from an XML file and generated using the *rphyloxml* and *ape* R packages, respectively.

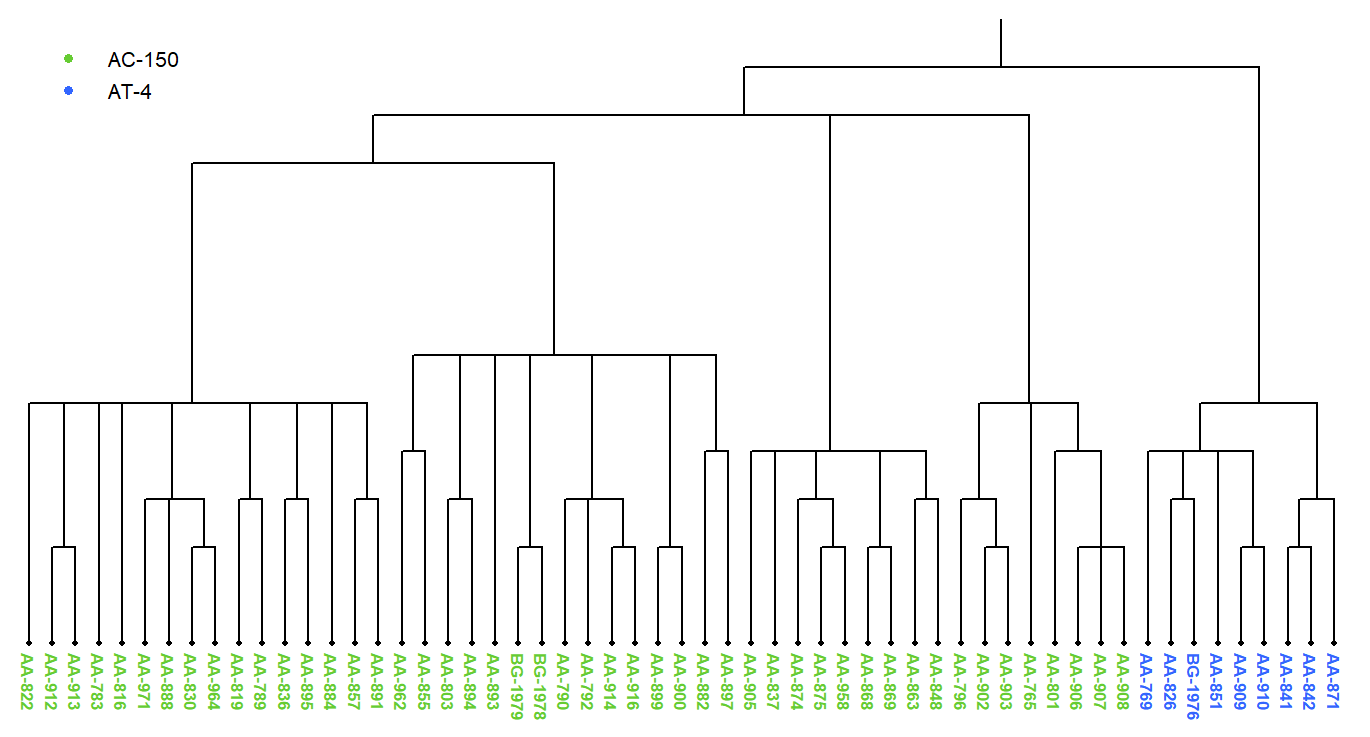

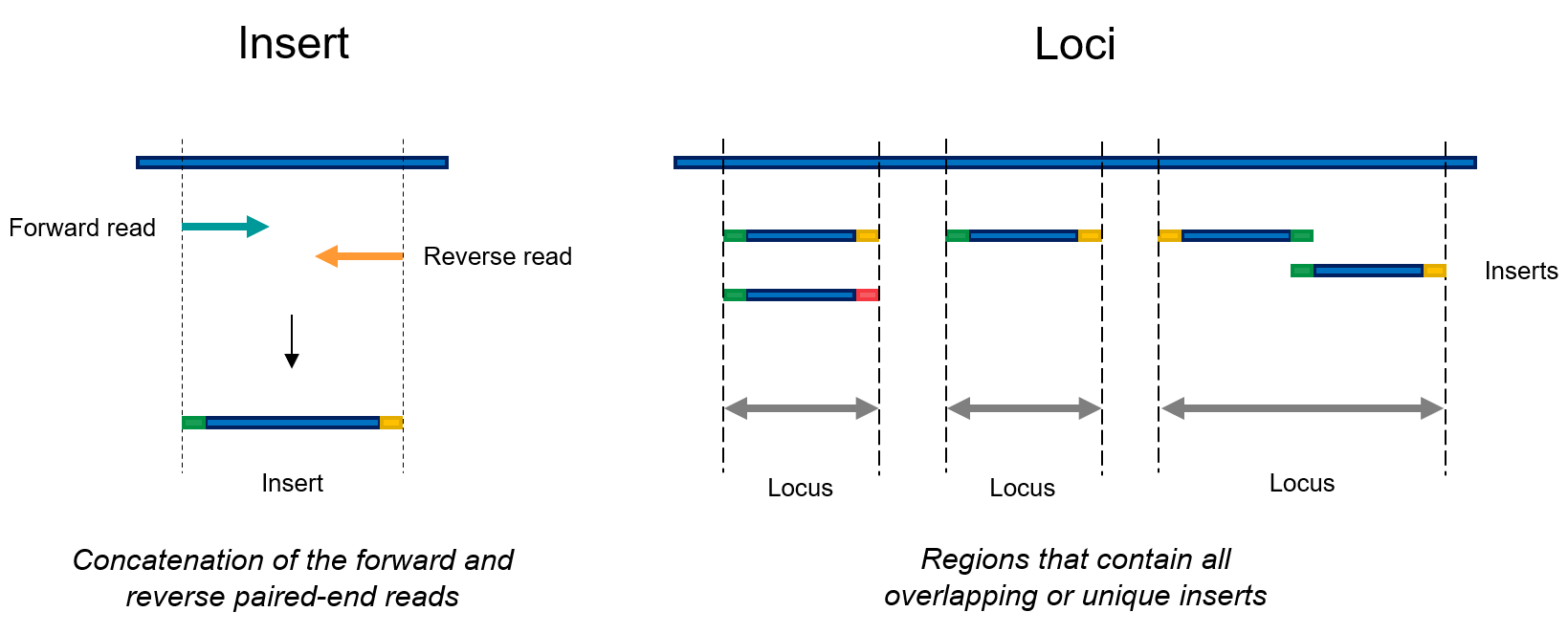
**Supplementary Figure 2.** Visual representation of inserts and loci as defined in the context of this study.

**Supplementary Results**

**Supplementary Table 1.** Genome coverage (%) and mean depth of coverage (X) for each CREAM library.

|  | **Genome coverage (%)** | | | **Mean depth of coverage (X)** | | |
| --- | --- | --- | --- | --- | --- | --- |
|  | ***Msp*I** | ***Hpa*II** | **Overall** | ***Msp*I** | ***Hpa*II** | **Overall** |
| **Mean** | 0.488 | 0.286 | 0.387 | 76.654 | 114.147 | 95.401 |
| **Max** | 0.885 | 0.629 | 0.885 | 164.033 | 231.384 | 231.384 |
| **Min** | 0.206 | 0.088 | 0.088 | 17.694 | 21.300 | 17.694 |

**Supplementary Figure 3.** Density of genes and the density of captured loci with the CREAM approach on the chromosomes of the cs10 cannabis reference genome in bins of 500 kb. Spearman’s rank correlation coefficient = 0.6913.

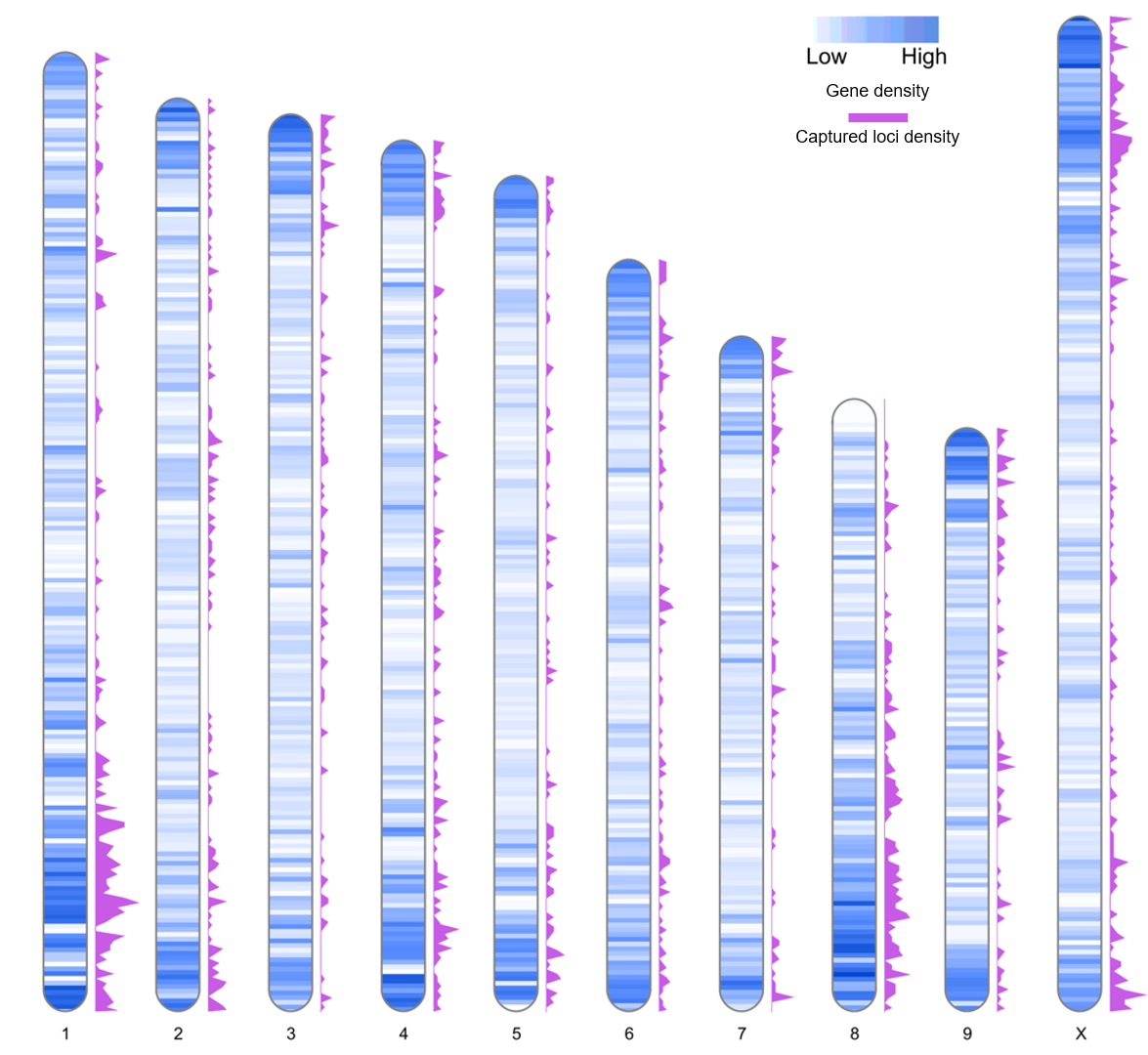
